# Expression of ERV3-1 in Leukocytes of Acute Myelogenous Leukemia Patients

**DOI:** 10.1101/2020.09.16.300905

**Authors:** So Nakagawa, Masaharu Kawashima, Yuji Miyatake, Kai Kudo, Ryutaro Kotaki, Kiyoshi Ando, Ai Kotani

## Abstract

Acute myelogenous leukemia (AML) is one of the major hematological malignancies. In the human genome, several have been found to originate from retroviruses, and some of which are involved in progression of various cancers. Hence, to investigate whether retroviral-like genes are associated with the development of AML, we conducted a transcriptome sequencing analysis of 12 retroviral-like genes of 150 AML patients using The Cancer Genome Atlas database. We found high expression of ERV3-1, an envelope gene of endogenous retrovirus group 3 member 1. In particular, blood and bone marrow cells of the myeloid lineage in AML patients, exhibited higher expression of ERV3-1 than those of the monocytic AML lineage. We also examined the protein expression of ERV3-1 by immunohistochemical analysis and found expression of ERV3-1 protein in 7 out of 12 AML patients, with a particular concentration observed at the membrane of some leukemic cells. Transcriptome analysis further suggested that upregulated ERV3-1 expression may be associated with chromosome 8 trisomy as anomaly was found to be more common among the high expression group compared to the low expression group. However, this finding was not corroborated by the immunohistochemical data. This discrepancy may have been caused, in part, by the small number of samples analyzed in this study. Although the precise associated molecular mechanisms remain unclear, our results suggest that ERV3-1 may be involved in AML development.

**Highlights:**

- Expression of 12 retroviral-like genes in the human genome were analyzed using transcriptome data of 150 acute myelogenous leukemia (AML) patients.
- ERV3-1, an envelope gene of endogenous retrovirus group 3 member 1, was found to uniquely show high expression level.
- Morphologic characteristics and chromosomal abnormalities are found to be related with the expression of ERV3-1.

## 1. Introduction

Approximately 8% of the human genome corresponds to retroviral origins (Lander et al. 2001). These areas of the genome are referred to as long terminal repeat (LTR) retrotransposons, many of which correspond to human endogenous retroviruses (HERVs). HERVs originally derived from retroviruses that infected germline cells of the host species. Therefore, HERVs contain retroviral genetic elements including cis-regulatory regions (LTRs) in their 5’ and 3’ terminals, as well as several coding sequences: gag, protease, polymerase, and envelope. The structures of other LTR retrotransposons are similar to that of HERV except for the absence of an envelope gene. Generally, HERVs are incapable of generating infectious virions that can competently replicate in human cells due to the accumulation of multiple mutations during evolution (Tönjes et al. 1999). Therefore, such retroviral sequences are believed to be “junk” DNA in the human genome. However, many recent studies showed that certain sequences, similar to those of retroviruses, have obtained new functions in the hosts.

In the human genome, at least 12 retroviral-like genes are annotated in the GRCh38 assembly provided by National Center for Biotechnology Information (NCBI): *ARC, ASPRV1/SASpase, ERV3-1, ERVK13-1, ERVH48-1/Suppressyn, ERVMER34-1/HEMO, ERVV-1, ERVV-2, ERVW-1/Syncytin-1, ERVFRD-1/Syncytin-2, PEG10/SIRH1, PEG11/RTL1/SIRH2, RTL4/ ZCCHC16/SIRH11* and *SIRH7/LDOC1. ERVW-1/Syncytin-1* and *ERVFRD-1/Syncytin-2* are the most well studied retroviral-like genes corresponding to retroviral envelope genes (Mi et al. 2000, Blaise et al. 2003), both of which are involved in human placenta development. Specifically, these genes are associated with cell-cell fusion and immunosuppression, both of which function are quite similar to those operated by envelope proteins of retroviruses (Kim et al. 2004). Those molecular functions may be also related to cancer progression. Indeed, Syncytin-1 and Syncytin-2 are reported to be involved in cancer development (Larsen et al. 2009). In addition, an LTR retrotransposon-derived PEG10/Sirh1 that is similar to a gag-pro-like gene is involved in placenta development (Ono et al. 2006), as well as in the progression of various cancers including pancreatic carcinoma, breast cancer, prostate cancer, gallbladder carcinoma, thyroid cancer, oral squamous cell carcinoma, colon cancer, enchondromas, and B-cell chronic lymphocytic leukemia (reviewed in Xie et al. 2018). Indeed, these retroviral-like genes originate from viruses making their unexpected expression potentially harmful to humans (Gonzalez-Cao et al. 2016).

Acute myelogenous leukemia (AML) is one of the major hematological malignancies, characterized by overproduction of myeloid progenitor cells in the bone marrow, which then rapidly migrates to the blood, and in some cases, can spread to other organs, such as liver and spleen. AML is associated with curative rates of 35 to 40% in patients aged < 60 years (Do□hner et al. 2010); however, the number of AML patients increase with age, and 70% of patients ≥ 65 years die of the disease within a year, despite treatment (Meyers et al. 2013). The French-American-British (FAB) classification system is a standard classification of AML patients that are divided into eight different subtypes (M0 through M7) based on morphologic characteristics (Bennett et al. 1976): undifferentiated acute myeloblastic leukemia (M0), acute myeloblastic leukemia with minimal maturation (M1), acute myeloblastic leukemia with maturation (M2), acute promyelocytic leukemia (M3), acute myelomonocytic leukemia (M4), acute monocytic leukemia (M5), acute erythroid leukemia (M6), and acute megakaryoblastic leukemia (M7).

Although numerous studies suggested relationships between HERVs and leukemia including AML (Depil et al. 2002; Chen et al. 2013; Bergallo et al. 2017; Cuellar et al. 2017; Deniz et al. 2020), details regarding the roles of retroviral-like genes in AML remain unclear, particularly as they pertain to the different AML subtypes. Therefore, in this study, we evaluated expression of retroviral-like genes in leukocytes of AML patients that are potentially harmful to AML. To this end, we first examined RNA-seq data obtained from 150 AML patients that were downloaded from The Cancer Genome Atlas (TCGA) database (https://www.cancer.gov/tcga). We then screened the expression of the abovementioned 12 retroviral-like genes and statistically examined the relationship between the expression levels and FAB subtypes, with exception of M6 and M7 cases, as they are relatively rare in AML (< 5%) (Bennett et al. 1976). We further validated the protein expression of the highly expressed retroviral-like gene in the leukemic cells obtained from AML patients by immunostaining and investigated whether the gene could be related to the progress of AML.

## 2. Materials and Methods

### 2.1 Ethics

This study was approved by the Institutional Review Board of Tokai University School of Medicine, of which protocol numbers are 15-I-26, 18-I-08 and 19-R-323 for immunohistochemistry and clinical sequencing data analyses of AML patients. Informed consent was provided according to the Helsinki Declaration in the Tokai University Hospital.

### 2.2 Cancer genome data analysis

Sequence and annotation data of the human genome GRCh38 was downloaded from the Illumina iGenomes (https://support.illumina.com/sequencing/sequencing_software/igenome.html). We also obtained RNA-seq data and clinical record data for 150 AML patients from TCGA-LAML database (https://portal.gdc.cancer.gov/projects/TCGA-LAML), which are summarized in the Supplementary data (Table S1 – S3). In this study, we used the RNA-seq data of which sequences are mapped to the human genome GRCh38 (BAM files) using STAR 2 (Dobin et al. 2013) provided by TCGA-LAML. We counted the mapped reads based on the gene annotation, and computed expression scores of TPM (transcripts per million) using StringTie2 version 2.0.6 (Kovaka et al. 2019). We extracted the TPM scores of 12 retroviral-like genes: *ARC, ASPRV1/SASpase, ERV3-1, ERVK13-1, ERVH48-1/Suppressyn, ERVMER34-1/HEMO, ERVV-1, ERVV-2, ERVW-1/Syncytin-1, ERVFRD-1/Syncytin-2, PEG10/SIRH1, PEG11/RTL1/SIRH2, RTL4/ZCCHC16/SIRH11*, and *SIRH7/LDOC1*, which are also summarized in the Supplementary data (Table S1). The TPM scores were log-transformed as follows: log_2_(TPM+1). Using the log-transformed TPM scores, we generated a heatmap of 12 retroviral-like genes using the heatmap.2 program in the gplots package of R (https://github.com/talgalili/gplots).

### 2.3 Statistical analysis

Normal variables were assessed by Fisher’s exact test. Continuous variables were assessed by Mann-Whitney *U* test or Kruskal-Wallis test for two or multiple groups, respectively. Data are presented as the mean ± standard deviation (SD). A *P* value < 0.05 was considered statistically significant.

### 2.4 Immunohistochemistry

To confirm ERV3-1 protein expression in AML, immunohistochemical (IHC) staining of ERV3-1 was performed on 12 cases of AML patients using paraffin-embedded bone marrow clot sections at Tokai University School of Medicine. Paraffin-embedded tissue sections were stained with hematoxylin-eosin. For immunostaining, an anti-human ERV3 antibody (rabbit polyclonal clone; Santa Cruz Biotechnology, CA), as a primary antibody, and anti-rabbit peroxidase histofine simple stain kit (Nichirei, Tokyo), as a secondary antibody, were used. The immunostaining tissue slides were observed by Olympus BX 63 microscope and cellSens software.

## 3. Results

We first examined the expression level of ERV3-1 from 150 RNA-seq data of blood and bone marrow of AML patients obtained from the TCGA database as summarized in the Supplementary data (Table S1). All sequencing reads were mapped to the human genome (GRCh38). Based on the mapped results, we measured the expression levels of all genes using the human genome annotation. We then compared the expression levels of 12 retroviral-like genes described in the Materials and Methods section. Figure 1 shows a heatmap of the expression in 150 samples measured by log-transformed TPM (transcripts per million) scores (see Materials and Methods). ERV3-1 was found to exhibit higher expression compared to eleven of the other retroviral-like genes. Indeed, the average and median TPM scores of ERV3-1 were 46.7 and 39.5, whereas those of the others were 2.4 and 0.2, respectively. We also examined the expression level of ERV3-1 in the GTEx database (https://www.gtexportal.org/), which collects various RNA-seq data from healthy people, and found that the median TPM score of whole blood is 4.6, and the highest expression score (27.3) was observed in adrenal gland. These results further indicate an upregulated expression of ERV3-1 in blood-bone marrow of AML patients.

**Figure 1.**
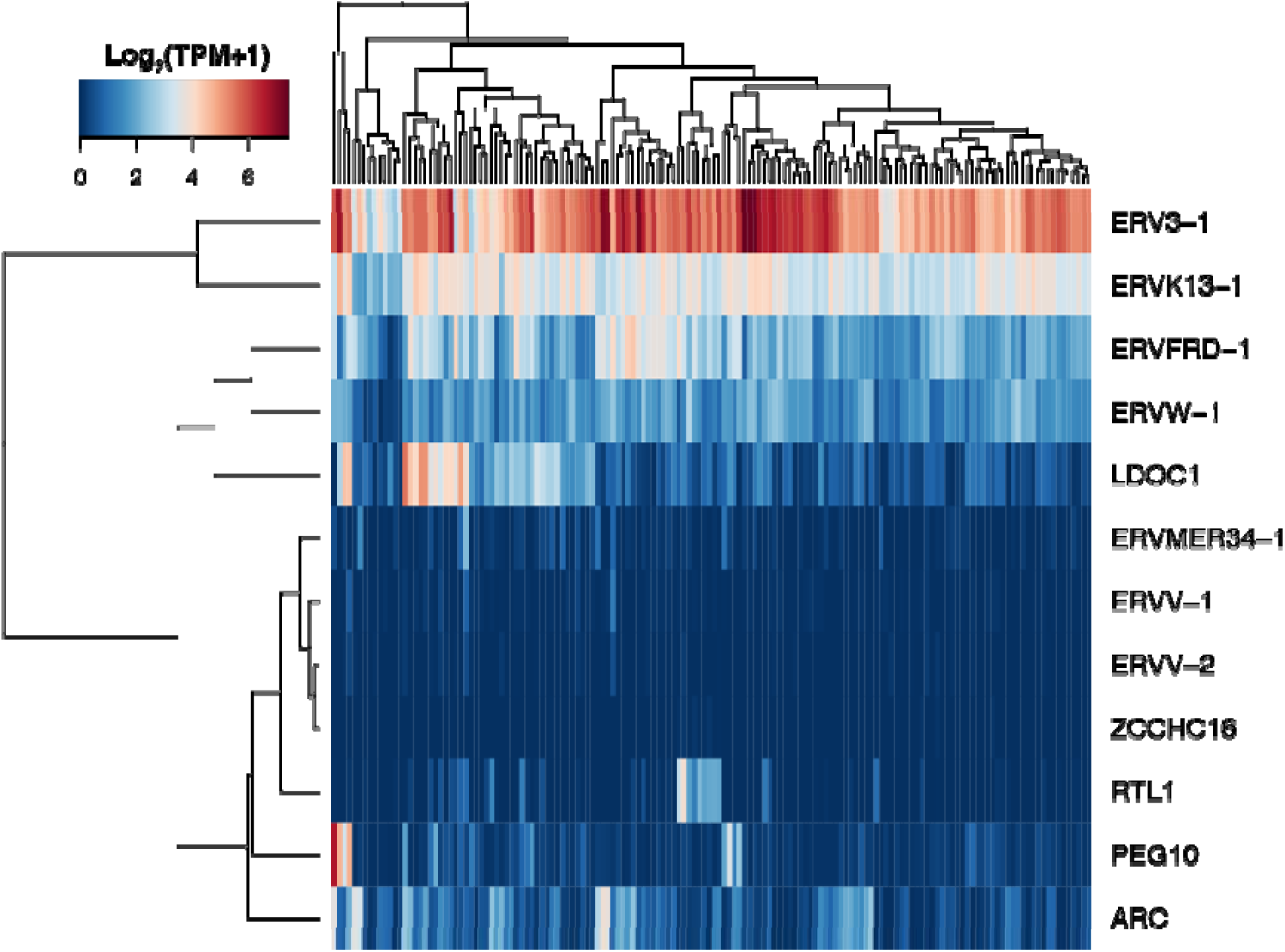
Transcriptome analysis of expression levels of retroviral-like genes. Heatmap of 11 retroviral-like genes 150 RNA-seq data is shown. Log-transformed TPM scores are used to compare the mRNA expression. Color in red or blue indicates the high or low expression levels, respectively.

We then evaluated relationship between ERV3-1 expression and clinical data, such as age, gender, cytogenetic risk, white cell count, and French-American-British (FAB) classification, as summarized in the Supplementary data (Table S2). We selected patients in the upper 20 and lower 20 percentiles of ERV3-1 expression (designated as the ERV3-1 high and low groups, respectively). In total, 60 patients were analyzed, the results for which are shown in Table 1 and the Supplementary data (Table S3). We found that ERV3-1 expression was not associated with age, gender, or white blood cell count, using the Mann-Whitney U test. Meanwhile, the cytogenetic risk is found to differ between the ERV3-1 high and low groups (*P* = 0.016). Moreover, the expression of ERV3-1 in AML FAB M0-M3 (myeloid phenotype) was higher than that of FAB M4-M5 (monocytic phenotype) (*P* < 0.001, Table 1 and Figure 2A). We then confirmed these observations using the whole 150 TCGA-LAML cases. All clinical data, excluding FAB classification, were not statistically associated with ERV3-1 expression (Figure S1). Hence, only FAB classification was statistically associated (*P* < 0.001, Figure 2B). Collectively, our transcriptome data analysis suggests that the blood and bone marrow of myeloid phenotype (FAB M0-M3) AML patients show higher expression levels of ERV3-1 than those of monocytic phenotype (FAB M4-M5).

**Table 1.** Characteristics of the TCGA-LAML patients in both ERV High (>20%) and low (<20%) groups.

| Patients | Total (n=60) | ERV3 High (n=30) | ERV3 low (n=30) | *P* |
| --- | --- | --- | --- | --- |
| Age: range (median) | 21-81 (56.5) | 21-81 (56.5) | 31-81 (56.5) | 0.706 |
| Gender |  |  |  |  |
| Male | 32 | 17 | 15 | 0.796 |
| Female | 28 | 13 | 15 |  |
| FAB classification |  |  |  |  |
| M0 | 7 | 5 | 2 | <0.001 |
| M1 | 18 | 11 | 7 |  |
| M2 | 15 | 9 | 6 |  |
| M3 | 4 | 4 | 0 |  |
| M4 | 10 | 1 | 9 |  |
| M5 | 6 | 0 | 6 |  |
| Cytogenetic risk |  |  |  |  |
| Favorable | 12 | 8 | 4 | 0.016 |
| Intermediate | 31 | 10 | 21 |  |
| Poor | 15 | 11 | 4 |  |
| Unknown | 2 | 1 | 1 |  |
| FAB classification |  |  |  |  |
| M0-M3 | 44 | 29 | 15 | <0.001 |
| M4-M5 | 16 | 1 | 15 |  |
| Diagnostic WBC:<br>range (median) | 1-58.5 (18) | 1-46 (12) | 1-58.5 (28.5) | 0.265 |
| ERV3-1 expression:<br>range (median) | 2.50-7.38 (5.36) | 6.13-7.38 (6.44) | 2.50-4.58 (4.02) | <0.001 |

WBC: white blood cell count

**Figure 2.**
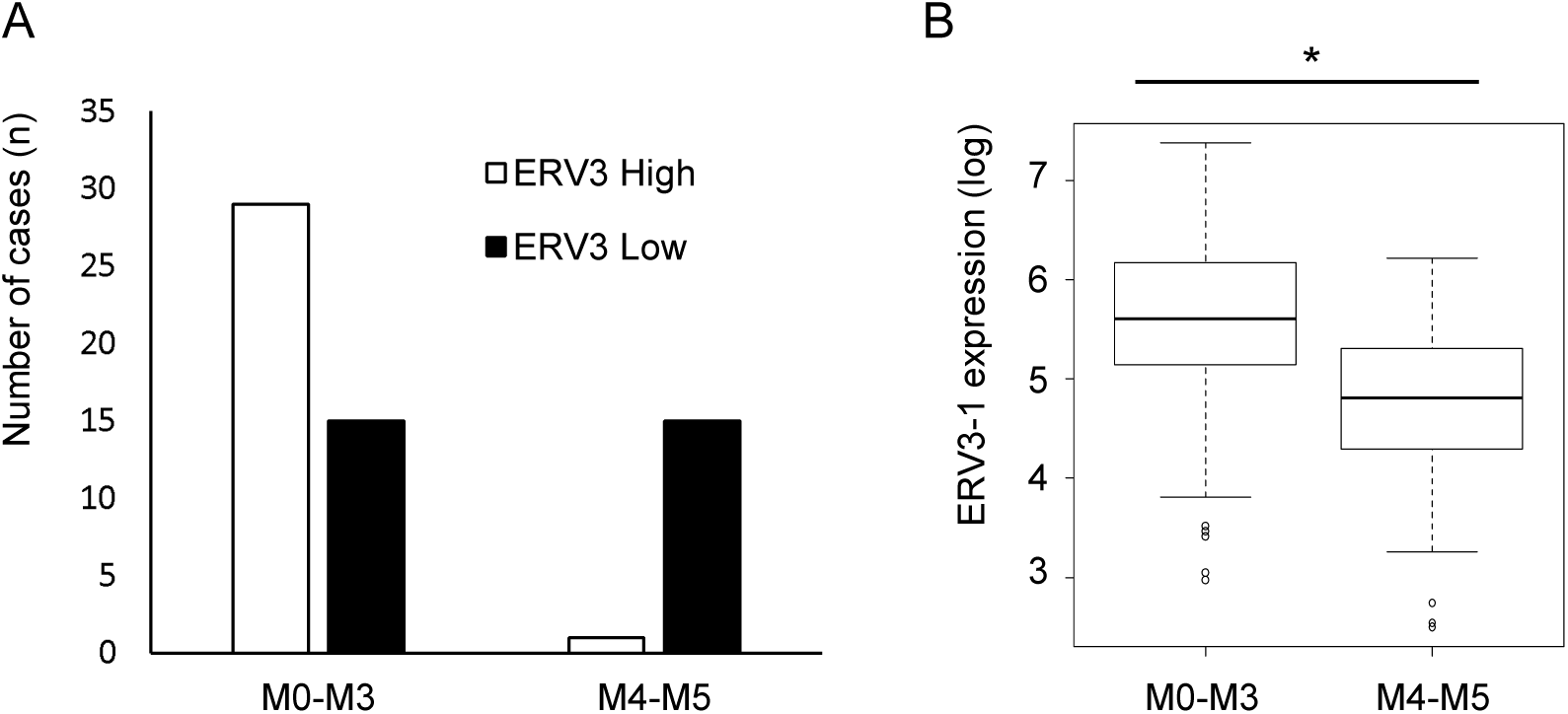
ERV3-1 expression of AML M0-M3 is higher than that of M4-M5. (A) Case distribution of FAB M0-M3 or M4-M5 in both ERV3-1 High (n= 30) and low groups (n=30) are shown. (B) Comparison of ERV3-1 expression of both AMLM0-M3 (n = 102) and M4-M5 (n = 44) in all cases of the TCGA data. Statistical analysis was assessed by Mann-Whitney U test. The boxes denote the median, and the first and third quartile. The upper and lower whiskers represent the 90% and 10%, respectively. *P < 0.001.

To examine the protein expression of ERV3-1 in bone marrow from AML patients, we conducted an immunohistochemical analysis for 12 AML patients at the Tokai University School of Medicine in Japan. Patients’ characteristics are summarized in Table 2. A previous study reported the expression of ERV3, including ERV3-1, in U-937 cells, which are one of AML cell lines classified as monocytic phenotype of AML (Larsson et al. 1996). Thus, we selected AML patients shown monocytic component classified as FAB M4-M5. In more than half of the cases (7/12), expression of ERV3-1 was detected, and, in particular, ERV3-1 was expressed at some of the leukemic cell membrane (Figure 3). The results clearly suggest that ERV3-1 RNA in blood-bone marrow of AML patients was translated and expressed as protein. Moreover, considering that our transcriptome analysis revealed low expression of ERV3-1 in M4-M5 group compared to M0-M3 group in the TCGA-LAML data (Figure 2), most M0-M3 probably cases likely contain ERV3-1 protein in tumor cells as well.

**Table 2.** Characteristics of AMLM4-M5 patients in in our immunostaining analysis

| Patients | Age at diagnosis | gender | FAB | Biopsy Status | Diagnostic White cell count | Cytogenetic | Cytogenetic risk | ERV3-1 expression |
| --- | --- | --- | --- | --- | --- | --- | --- | --- |
| 1 | 79 | male | M4 | Primary | 2000 | 45, X, -Y [20/20] | Intermediate | + |
| 2 | 78 | male | M4 | Primary | 27700 | 46,XY [20/20] | Intermediate | ++ |
| 3 | 60 | female | M4 | Primary | 17900 | 45, XX, -7 [20/20] | poor | – |
| 4 | 68 | female | M4 | Primary | 34700 | 46, XX,<br>t(7;11)(p15;p15)<br>[20/20] | Intermediate | – |
| 5 | 69 | male | M4 | Primary | 1500 | 46,XY,der(7)(q31),<br>Inv(16)(p13.1q22)<br>[20/20] | favorable | + |
| 6 | 53 | male | M5 | Relapse | 48600 | 46,XY [20/20] | Intermediate | ++ |
| 7 | 70 | male | M5 | Primary | 33800 | 46,XY, inv(9)(p12q13)<br>[10/20] | Intermediate | + |
| 8 | 65 | male | M5 | Primary | 61000 | 47, XY, -8, +i(8)(q10)<br>×2 [20/20] | Intermediate | – |
| 9 | 64 | male | M5 | Primary | 50500 | 49,XY,+5,add(7)(q32),<br>+8,+mar [8/20]<br>/50,idem,+20[3/20] | Poor | ++ |
| 10 | 58 | male | M5 | Primary | 17400 | 47,XY,+8 [18/20] | Intermediate | – |
| 11 | 56 | male | M5 | Primary | 169800 | 46,XY [20/20] | Intermediate | + |
| 12 | 71 | male | M5 | Primary | 5700 | 46,XY [20/20] | Intermediate | – |

**Figure 3.**
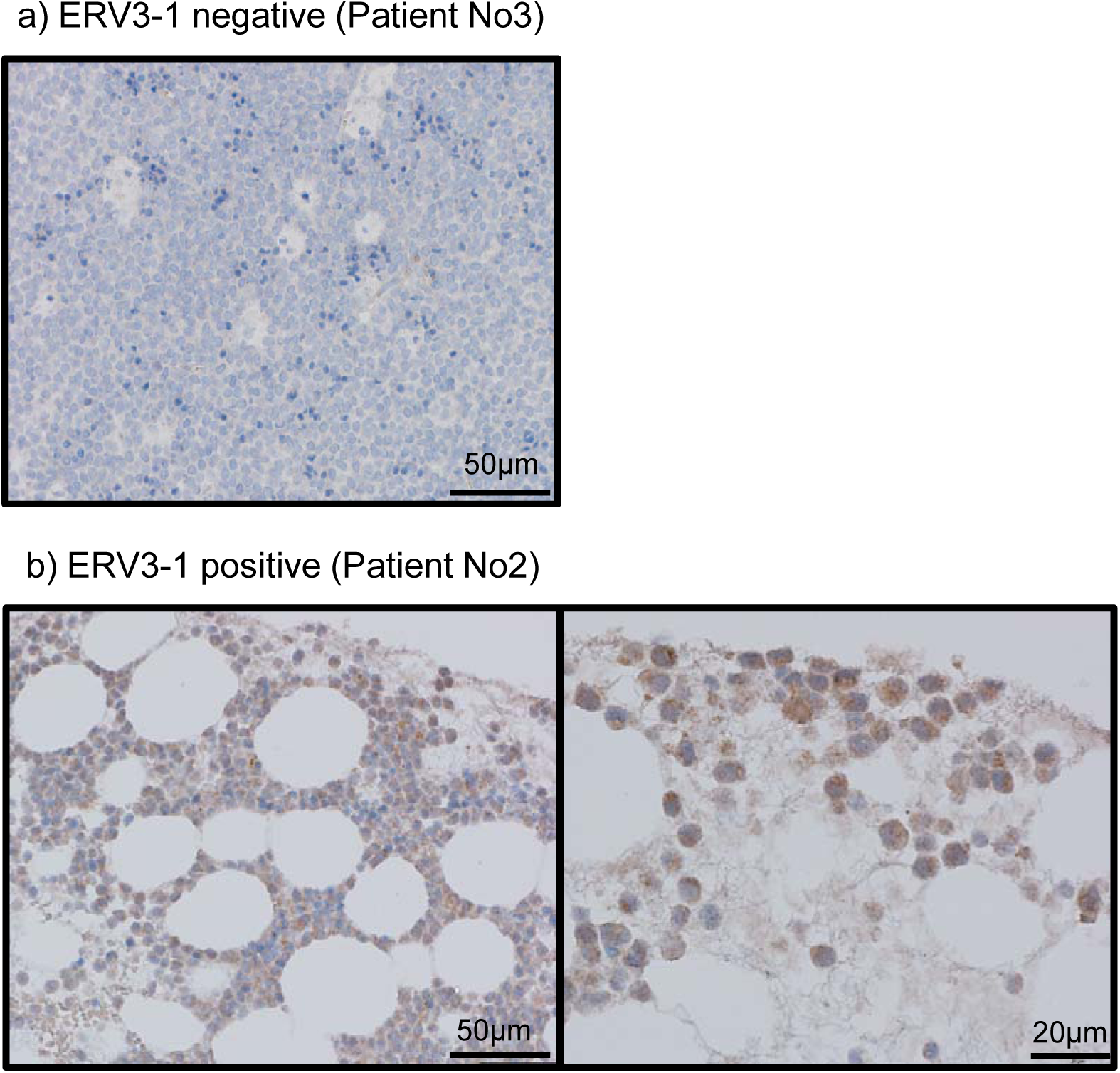
More than half of AML M4-M5 patients express ERV3-1 protein in tumor cells. Immunohistochemical staining of ERV3-1 was performed using bone marrow of patient samples. Tumor cells were occupied in the majority of bone marrow tissue. Representative of ERV3-1 (A) negative and (B) positive patients (left: low power field, right: high power field) corresponding to Table 2 are shown.

We also evaluated association between ERV3-1 expression and chromosomal abnormalities and genetic mutations, which are considered to be involved in AML progression (Short et al. 2018). Specifically, chromosomal abnormalities, such as translocation chromosomes t (15;17) and t (8;21), and trisomy of chromosome 8, are reportedly associated with AML (Vickers et al. 2000). We, therefore, focused on these anomalies in our analysis. Results show that trisomy 8 was more common in the ERV3-1 high group compared to those of the low group (Table S1, *P* = 0.0232). In our immunostaining analysis, however, the prevalence of trisomy 8 has not been a clear difference in both ERV3-1 positive (only Patient 9) and negative (Patient 10) cases (Table 2). We also evaluated three major mutations that related with AML: fms-related tyrosine kinase 3 (FLT3), isocitrate dehydrogenase 1 (IDH1), and nucleophosmin 1 (NPM1); however, no significant associations were detected between these mutations and ERV3-1 expression, as shown in the Supplementary data (Table S4).

## 4. Discussion

Although many retroviral-like genes have been shown to be related to cancer development (Gonzalez-Cao et al. 2016), here we specifically found that ERV3-1 shows exclusively a high expression level in blood and bone marrow of all of AML patients using TCGA database (Figure 1). We also confirmed that ERV3-1 protein was detected in more than half of AML M4-M5 patients (7/12) (Figure 3). Although we have not examined the protein expression of ERV3-1 in blood-bone marrow of AML M0-M3, we found that mRNA expression level is higher in AML M0-M3 than in AML M4-M5 (Figure 2) suggesting that patients of AML M0-M3 may express the ERV3-1 protein as well. Those results indicate that ERV3-1 protein as well as mRNAs may be expressed in blood-bone marrow of most of AML patients.

ERV3-1 is an envelope gene of the endogenous retrovirus group 3 member 1, which belongs to the HERV-R family. It is known that retroviral envelope gene is involved in various biological processes, including infection and immunosuppression. Indeed, ERV3-1 was reportedly expressed in placenta (Venables et al. 1995; Lin et al. 2000; Blaise et al. 2007) and in colorectal cancers (Lee et al. 2014). Although ERV3-1 lost its fusogenic activity (Blaise et al. 2007), it contains an immunosuppressive region in the transmembrane domain, termed p15E, of C-type retroviruses, suggesting that ERV3-1 may serve to suppress immune response (Venables et al. 1995). Indeed, immunosuppressive region of another retroviral envelope-derived gene, syncytin-2, supports the injection of MCA205 mouse fibrosarcoma cell line in mice (Mangeney et al. 2007). Therefore, immunosuppressive activity of ERV3-1 could potentially be related to the progress of AML.

AML forms an immunosuppressive microenvironment by increasing the number of myeloid-derived suppressor cells in the peripheral blood, as well as regulatory T cells in both the peripheral blood and bone marrow (Beyar-Katz et al. 2018). In fact, allogeneic hematopoietic cell transplantation, one of the T-cell based immunotherapy, is the most effective in post-remission therapy, and is commonly used for AML treatment (Koreth et al. 2009). AML cell spontaneously fused with murine macrophages, endothelial, and dendritic cells, which may lead to dissemination of the disease (Martin-Padura et al. 2012). This observation suggests that immunosuppressive function of ERV3-1 might be involved in AML progression.

Although 150 cases show high ERV3-1 mRNA expression levels (Figure 1), we were unable to confirm the protein expression of ERV3-1 in 5 of 12 cases (Figure 3). We were also unable to identify an association of ERV3-1 expression with chromosomal abnormalities and genetic mutations (Table S1). These results might suggest that ERV3-1 is not an essential factor in AML development, but rather plays a supportive role. Therefore, the factor that affects ERV3-1 expression of AML, as well as the role of ERV3-1 in AML, should be further investigated. Moreover, considering that many viral-derived sequences have been described in eukaryote genomes that have not yet been annotated in the genome database (Nakagawa and Takahashi 2016; Pertea et al. 2018; Kryukov et al. 2019) and that these viral-derived genes are dynamically altered during evolution (Imakawa et al. 2015; Imakawa and Nakagawa 2017; Pastuzyn et al. 2018). Therefore, not only ERV3-1 but also other unknown viral-derived genes could be also involved in the progress of AML.

## Abbreviations

AML: acute myelogenous leukemia
ERV: endogenous retrovirus
FAB: French-American-British
LTR: long terminal repeat
FAB: French-American-British Classification
TCGA: The Cancer Genome Atlas
TPM: transcripts per million

## Author’s contributions

S.N. and A.K. conceived the study idea. S.N. and M.K. conducted the data analysis. Y.M., K.K., R.K. and A.K. conducted experiments. M.K., K.A. and A.K. interpreted the data. S.N., M.K. and A.K. wrote the manuscript. All authors read and approved the final manuscript.

## Acknowledgement

The results shown here are in part based upon data generated by the TCGA Research Network: https://www.cancer.gov/tcga. Computations in this work were performed in part on the NIG supercomputer at ROIS National Institute of Genetics and SHIROKANE at Human Genome Center (the Univ. of Tokyo).

