## Supplementary figure for "Expression of ERV3-1 in Leukocytes of Acute Myelogenous Leukemia Patients"

### Slide 1
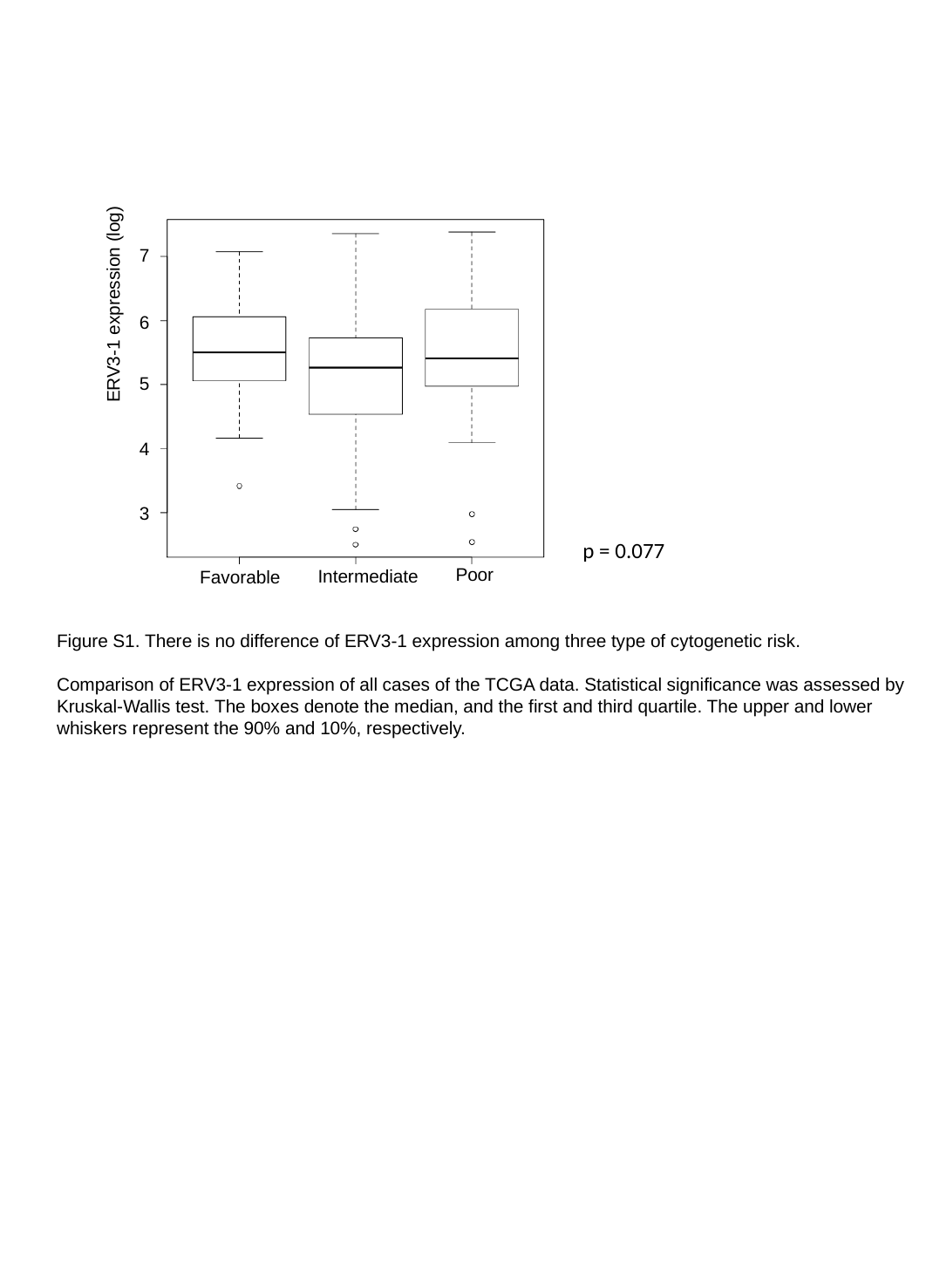

7
ERV3-1 expression (log)
6
5
4
3
p = 0.077
Poor
Intermediate
Favorable
Figure S1. There is no difference of ERV3-1 expression among three type of cytogenetic risk.
Comparison of ERV3-1 expression of all cases of the TCGA data. Statistical significance was assessed by Kruskal-Wallis test. The boxes denote the median, and the first and third quartile. The upper and lower whiskers represent the 90% and 10%, respectively.
